# EpicPCR-Directed Cultivation of a *Candidatus* Saccharibacteria Symbiont Reveals a Type IV Pili-dependent Epibiotic Lifestyle

**DOI:** 10.1101/2021.07.08.451036

**Authors:** Bingliang Xie, Jian Wang, Yong Nie, Dongwei Chen, Beiyu Hu, Xiaolei Wu, Wenbin Du

**Affiliations:** State Key Laboratory of Microbial Resources, Institute of Microbiology, Chinese Academy of Sciences, Beijing 100101, China; College of Engineering, Peking University, 100871 Beijing, China; College of Life Sciences, University of the Chinese Academy of Sciences, Beijing, 10049, China; Savaid Medical School, University of the Chinese Academy of Sciences, Beijing 100049, China; State Key Laboratory of Transducer Technology, Institute of Microbiology, Chinese Academy of Sciences, Beijing 100101, China

## Abstract

Candidate phyla radiations (CPR), accounting for a major microbial supergroup with remarkably small genomes and reduced sizes, are widely distributed yet mostly uncultured. Limited culture and its obligate reliance upon other bacteria hindered investigation of their lifestyles. In this work we isolated a CPR bacterium, TM7i, with its host *Leucobacter aridocollis* J1, by combination of Emulsion, Paired Isolation and Concatenation PCR (epicPCR) detection and filtrate co-culture. Genomic profiling of TM7 genomes and microscopic investigation of TM7i-J1 symbiosis suggest the conservation of type IV pili and a pili-dependent lifestyle of TM7. Further, we observed twitching motility of TM7i mediated by pili and its role played in the interaction with its host. Our results shed a light on the lifestyle about this enigmatic bacterial radiation, which may also be adopted by other CPR organisms. The epicPCR-directed isolation method underlines high efficiency of CPR bacteria isolation and thus may be used in other symbiotic or epibiotic microorganisms.

## Introduction

The majority of bacteria in the environment remains to be cultured^1^. This phenomenon is an obstacle for microbiology and hindered the understanding and exploration of massive microbial resources. Various methods have been introduced, including gel microdroplets^2^, microfluidic droplet arrays^3^, *In Situ* diffusion chamber^4^, iChip^5^, and SlipChip^6^, *etc*. These methods have revealed higher culturable biodiversity and enriched new bacterial groups. However, most of the classic strategies for microbial studies are based on “pure culture”, and are inapplicable to isolation of symbiotic species with obligate partner-dependent lifestyles.

Using culture-independent methods, microbial groups with ecological and evolutional importance have been discovered and expanded the tree of life^7^. Candidate phyla radiation (CPR) is one kind of these widely distributed but mostly uncultured groups, accounting for over 15% of total microbial biodiversity in the whole bacterial domain^8^. CPR is distinctive in its extremely small genome^8^, tiny cell size^9^, and epibiotic lifestyles which is merely understood^10, 11, 12^. Due to the lack of complete glycolysis, lipid biosynthesis, and various amino acid biosynthesis pathways, CPR bacteria are considered to be obligate epibionts based on genomic analysis and cultivation, which means they need a partner to accomplish their lifecycle^9, 13, 14^.

TM7, *Candidatus* Saccharibacteria^7^, is a CPR phylum widely distributed in many environments such as human oral cavity^15, 16^, groundwater^13^, rhizosphere soil^17^, rumen^8, 18^, *etc*. As the first and only confirmed epibiotic CPR, TM7 need actinobacteria as their hosts to accomplish a parasitic lifestyle^11, 14, 19, 20^. Insights to TM7 symbiosis would facilitate the exploitation of uncultured microbes and promote the understanding of minimal genomes for synthetic biology. Studies have shown that TM7 has an extremely streamlined genome, making it incapable of utilizing glycans and synthesizing membrane lipids^13^. Also, a rapid evolution occurred when *Schaalia odontolytica* XH001 was infected by TM7x^12^. The conservation of Type II Secretion System (T2SS) in the TM7 group indicates a T2SS-dependent symbiotic lifestyle^21^. These clues indicate that there is complex substance exchange between TM7 and its host. However, little is known about TM7’s lifecycle due to very few cultured representatives. The ultra-small size of TM7 (∼200 nm) hinders its visualization using conventional optical microscopy. Crucial questions arise: how TM7 cells move, select, and attach to their host; new efforts should be directed toward improving the isolation of TM7 symbionts from various environments other than the human oral cavity, as well as a better understanding of the biological mechanisms of its epibiotic lifestyle. Bacterial type IV pili (T4P) has been functioned to facilitate motility^22^, competence^23^, form aggregating organization^24^. Previous genomic studies indicate that T4P is enriched in CPR, including some TM7 representatives^25^, which may facilitate CPR symbiosis. However, the contribution of T4P in TM7’s lifecycle remains unknown.

There is still limited understanding of CPR as well as other obligate epibionts such as DPANN nanoarchaea, due to the lack of efficient methods for isolating these microbes. For isolation of obligate symbiotic species such as TM7, a targeted technique with high efficiency is required. TM7 has been enriched and cocultured from oral samples using streptomycin for selection^14^. However, during enrichment, TM7 symbionts suffer from intense competition with fast-growing antibiotic-resistant microbes and are likely to decline during passages. Recently, oral TM7 and its hosts are co-isolated using Fluorescence-Activated Cell Sorting (FACS) and fluorescence immunolabeling^26^. However, the immuno-targeted method is hard to explore TM7-host interaction at a systematic level and lacks flexibility when changing targets due to costly antibody production and purification. Besides, except for limited TM7 co-isolates from the oral cavity, no environmental-related TM7 symbiont has been isolated yet.

Genetic-based methods, such as PCR^6^ and 16S rRNA gene sequencing, can detect target CPR in a sample but can not reveal the symbiotic relationships in complex microbial communities. epicPCR (Emulsion, Paired Isolation and Concatenation PCR)^27^ is first introduced for linking functional genes with phylogenetic markers of single cells in a community by single-cell isolation and concatenation PCR in hydrogel emulsions. Recently, EpicPCR has been adopted to detect interspecies relationships with close spatial interaction, which can be co-isolated in droplet emulsions. EpicPCR detections of phage-bacteria^28^ infection and bacteria-protozoan^29^ endosymbionts have been demonstrated. According to these studies, we envision that epicPCR can also be used for detecting bacterial epibiotic relationships, especially CPR-host symbionts in a microbial community.

In this study, we introduced a new approach combining epicPCR and selective co-cultivation for targeted enriching and isolating CPR-host symbionts. Following this simple workflow, we successfully co-isolated TM7i with its host *Leucobacter aridocollis* J1 from a cicada exuviae sample. We found TM7i different from other oral TM7 in both phylogenetics and physiological interactions. Further, using electron microscopy and super-resolution microscopy, we observed T4P-dependent motility and interaction of TM7i with J1, which shed light on the symbiosis lifecycle of TM7.

## Results

### The epicPCR-directed CPR-host symbionts isolation approach

A simple workflow of epicPCR-directed cocultivation is introduced to dramatically improve the selectivity and efficiency for CPR-host symbiont discovery (Figure 1). First, we use epicPCR^30^ to detect epibiotic CPRs associated with the hosts in a microbial community (Fig. 1a). Briefly, we prepared bacterial suspensions containing CPR-host symbionts from environmental samples or enrichment cultures. We emulsified them into water-in-oil picoliter polyacrylamide droplets stochastically containing either a free-living cell, a CPR-affected host cell, or no cell. The droplets are solidified into hydrogel beads followed by lysozyme treatment for bacteria lysis. The beads are re-emulsified together with PCR mix for fusion PCR to generate concatenated amplicons link sequences of CPRs with their hosts. In this study, particularly, a fusion PCR is designed for TM7-host symbionts, yielding ∼800bp fused amplicons combining ∼300bp 16S rRNA gene sequences of TM7 and ∼500bp 16S rRNA gene sequences of their hosts. Finally, a bulk nested PCR is performed to add Illumina adapters and the TM7-host symbionts information is generated by next-generation sequencing.

**Figure 1.**
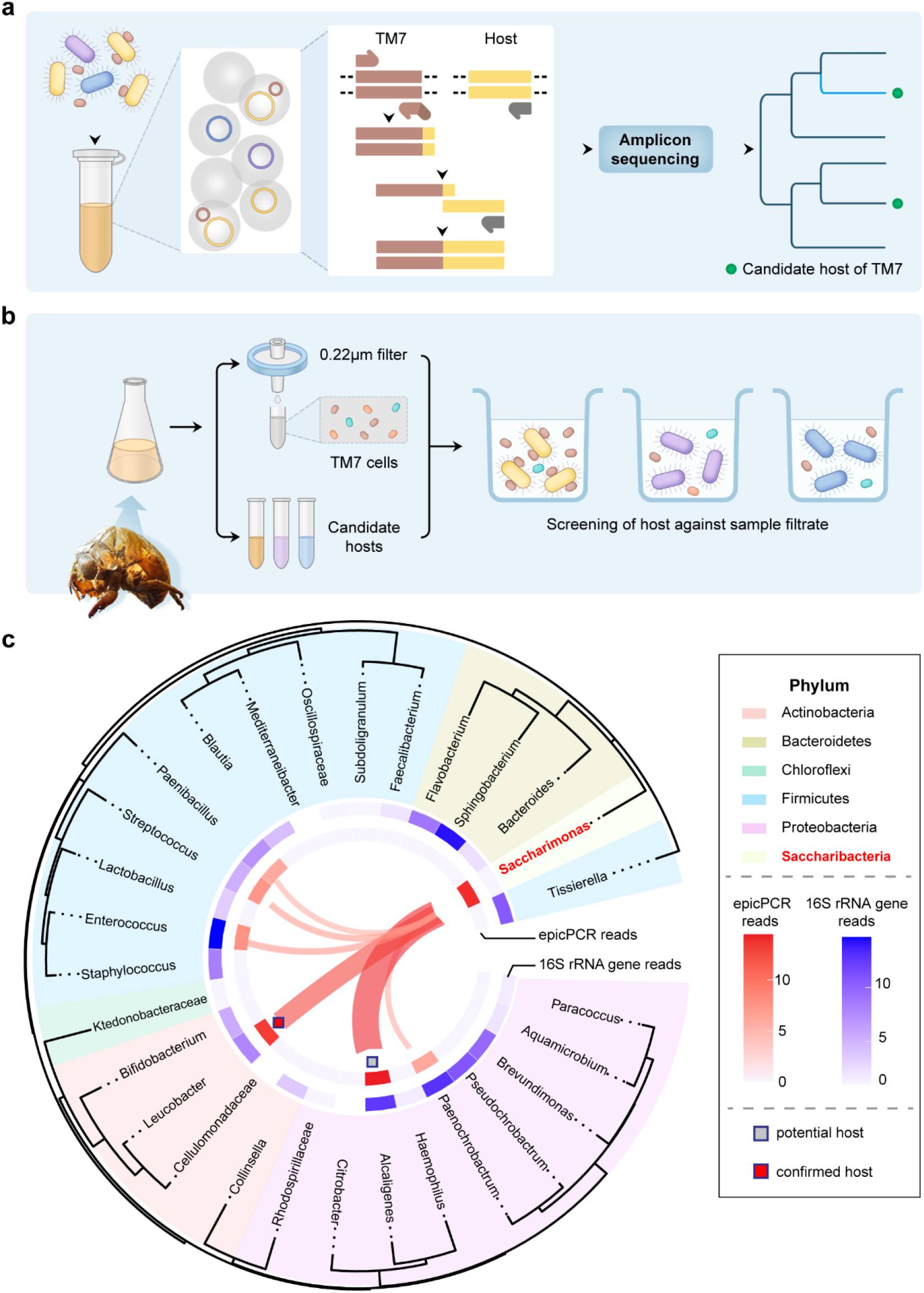
epicPCR-Directed isolation of *Candidatus* Saccharibacteria (TM7) and its symbiotic microbial hosts. **a**, epicPCR (Emulsion, Paired Isolation and Concatenation PCR) is used to identify the hosts of TM7 in an enrichment community. Single microbial cells, including host cells infected with TM7, are encapsulated in picoliter polyacrylamide emulsions. Fusion PCR and amplicon sequencing were performed to link 16S rRNA genes of hosts and TM7 to detect microbial symbionts. **b**, Guided by hosts information obtained by epicPCR, candidate hosts were screened against 0.22 μm filtrate of a cicada-related enrichment (which contains free TM7 cells) to obtain TM7-host symbionts. **c**, TM7-host relationships and relative abundance of the cicada exuviae aerobic enrichment community revealed by epicPCR. Community-level abundance (outer ring, blue) and epicPCR-revealed host abundance (inner ring, red) were provided.

Secondly, the information revealed by epicPCR allows us to quickly identify potential hosts in a complex microbial community to guide the selective isolation of TM7-host symbionts (Fig. 1b). We can directly isolate the putative hosts from the samples or obtain representative strains from commercially available resources. Then, small particles, including free TM7 cells, are separated from the microbial community suspensions by filtration using 0.22-μm filter membrane based on its ultrasmall size compare with other bacterial species^9, 14^. Finally, the putative hosts are screened against the filtrate by co-cultivation and PCR targeting the growth of TM7. This combined technique, including epicPCR and filtrate co-cultivation, significantly increased the efficiency and reduced uncertainty for TM7-host isolation.

### Validation of epicPCR targeting TM7 symbionts using an artificial sample

To validate TM7-host detection using epicPCR, we constructed an artificial TM7-host symbiont. *Escherichia coli* (*E. coli*) transformed with a plasmid containing TM7 16S rRNA gene sequence was used to simulate a TM7-*E. coli* symbiont, mimic the close contact between the host and TM7 (Supplementary Fig. 1a). The experiment consists of three representative samples: the TM7-positive community sample combines transformed *E. coli* and *Staphylococcus aureus* (*S. aureus*); the TM7-free community is composed of wild type *E. coli* mixed with *S. aureus*; the pure ‘TM7-host symbiont’ containing only transformed *E. coli* (Supplementary Fig. 1a). After performed epicPCR with the above three samples, the fusion PCR successfully detected the artificial TM7-host symbiont in the positive community and the pure TM7-host symbiont with 84.9% and 98.5% of total epicPCR reads, respectively. And the TM7-free community shows no nested PCR products and no fused amplicon, proving the high specificity of our TM7-host detection method (Supplementary Fig. 1b,c). Therefore, the primer set we designed can detect TM7-host symbiont from a mixture. Further, we can detect various epibiotic symbionts from the community with low contamination and high sensitivity by developing epicPCR primer sets targeting different CPRs.

### epicPCR-directed isolation of an environmental TM7 Symbiont

We detected TM7i microbial groups by lineage-specific PCR from an enriched sample from cicada exuviae. After continuous passages, we can stably detect the existence of TM7i, which indicates there are host-TM7i symbionts in the enriched sample (Fig. 2a). We also detected sequences of TM7i in 0.22-μm filtrate of the enrichment, proving that the TM7 in our sample also has representative ultrasmall cell size^3,8^. To test if epicPCR can reveal TM7-host symbionts in the enrichment, we performed epicPCR and 16S rRNA gene amplicon sequencing. The epicPCR indicates highly linkage between *Saccharimonas* gene sequences with *Alcaligenes* and *Leucobacter* (Fig 1c). Afterward, we successfully isolated *Alcaligenes* spp. And *Leucobacter aridocollis* J1 from the sample and cocultured them with 0.22-μm filtrate to see if they are the hosts of TM7i. Fortunately, TM7i is amplified with the J1 strain and preserved during multiple passages with J1 in broth and agar culture. (Fig. 2a). This indicates J1 is the host of the TM7i, and the sequencing clue provided by epicPCR can guide the cultivation of TM7-host Symbionts.

**Figure 2.**
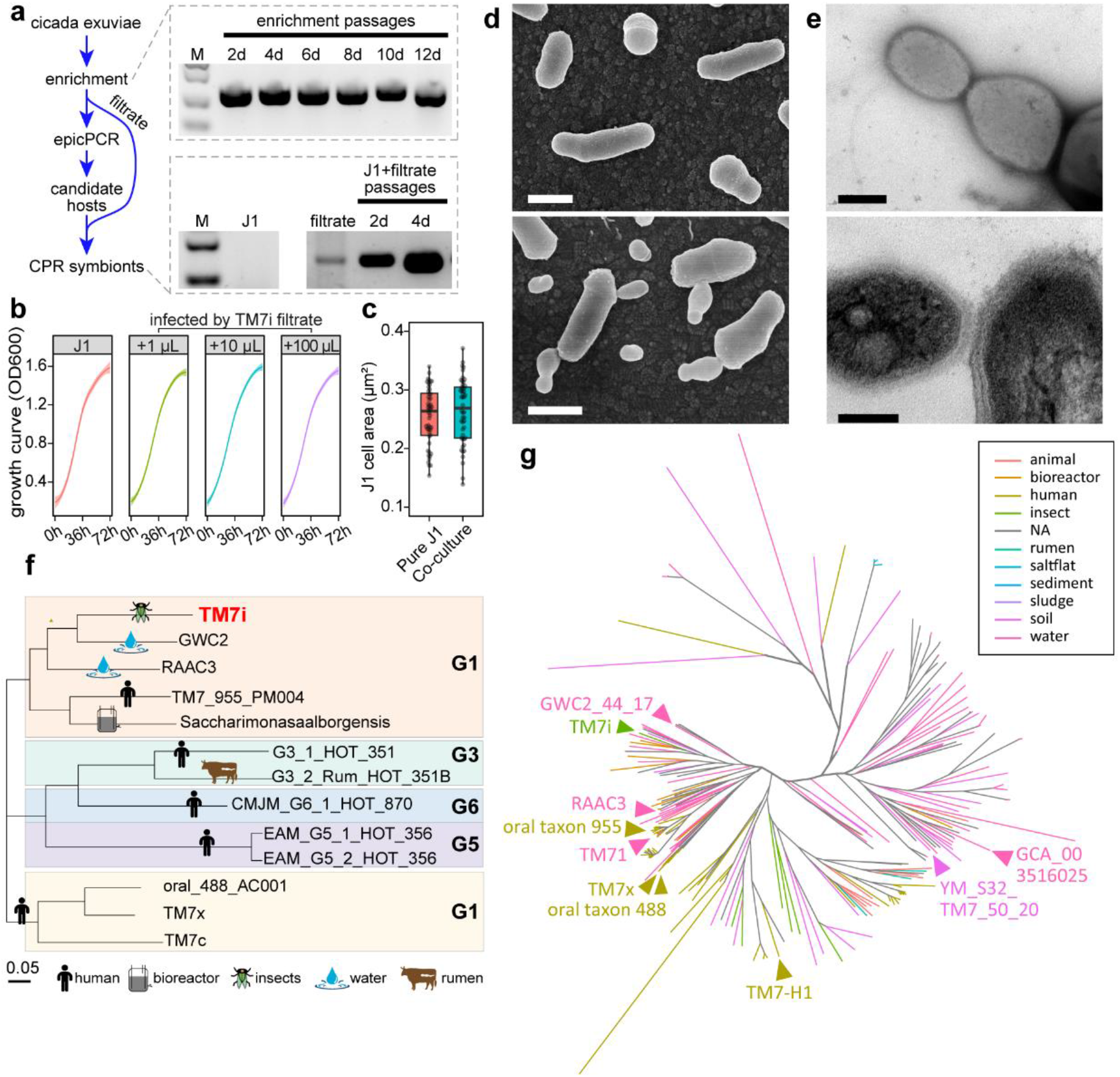
Cultivation of a novel *Candidatus* Saccharibacteria (TM7i) and its host *Leucobacter aridocollis* J1 (J1). **a**, Workflow of epic-PCR directed TM7 isolation in a cicada exuviae enrichment. Existence of TM7 symbiont in the enrichment confirmed by continuous passages (up right). Obtained TM7i-J1 symbiont was validated by multiple passages and PCR detection of TM7 (down right). **b**, Growth curve of pure J1 and infected J1 with different volume of filtrate containing TM7i. **c**, the cell size of J1 before and after co-culture. **d**, the shape of J1 did not change upon co-culture. scale bar =500nm. **e**, Transmission electron microscopy (TEM) reveal uneven binary fission of TM7i (up) and close adhesion of TM7i with J1 (bottom). scale bar=200nm. **f**,**g**, Phylogenetic tree of genome assemblies of TM7 of different habitats.

### Symbiosis of TM7i with *Leucobacter aridocollis* J1

TM7i, like other cultured TM7 representatives, relies exclusively on its host to grow and proliferate. But compared with cultured and studied TM7x, TM7i is different in three aspects: habitat of isolation, phylogenetic distance, and the symbiotic relationship. TM7i and its host J1 are isolated from a sample enriched from Cicada exuviae, known as traditional herbal medicine. To our knowledge, it is the first TM7-host symbiont isolated from an insect host. The occurrence of TM7 with cicada is rarely reported and not well studied. However, Cicadas live underground as nymphs for most of their lives, and their exuviae give supports to the growth of an unique microbial community associated with the forestry soil microbiota. Meanwhile, the forest soil is a rich reservoir of fungus and there is dynamic equilibrium among insect, fungi and actinobacteria, because fungi act as mutualists protecting insects from fungal infection^31, 32^. Therefore, we hypothesize that the cicada nymphs use their shell to recruited soil actinobacteria with antifungal activities, and actinobacteria further host the growth of epibiotic TM7 species. Genome-based phylogenetic analysis based on up-to-date bacterial core gene (UBCG)^33^ method reveals that TM7i belongs to another major clade which is distant from cultured oral TM7 isolates (Fig. 2f,g); this is in accordance with habitat-association of the evolutionary and diversification processes^34^. TM7x is the first TM7 isolate obtained from the oral sample, with *Schaalia odontolytica* XH001 as its host and isolated in anaerobic or microaerobic environment^14^. Comparison of 16S rRNA gene similarity shows TM7i is only related with TM7x at 93% similarity and is more distant from other cultivated TM7 strains (Supplementary Fig. 2).

We also found the symbiotic relationship between TM7i and its host is very different from previous studies. Recent studies have shown that the epiparasitic symbiosis of TM7x and its host XH001 triggered cellular stress response similar to oxygen-depletion stress, which resulted in morphological changes and negatively affected the growth of XH001^11^. Meanwhile, differet responses from various hosts of TM7x was observed^35^. In comparison, TM7i and J1 symbiont grow with aerobic conditions, and it seems that both the growth and morphology of J1 are not affected by epibiotic symbiosis with TM7i. The growth curves show that the optical density is identical for pure J1 culture and the TM7i-J1 symbiont (Fig. 2b). The images obtained by scanning electron microscopy (SEM) show that the shape and cell size of J1 have no significant difference after infection by TM7i (Fig. 2c, Fig. 2d). These results indicate that TM7i and TM7x treat differently to their host or their hosts have different ways of adapting to the symbiosis with TM7. However, more physiological and metabolic studies should be carried out to reveal the mechanism of symbiotic interaction between TM7i and J1.

One intriguing point is that large-scale ultra-thin slice TEM imaging showed no cell membrane integration between TM7i and the host was observed: the transparent cell wall of each cell separates them from each other (Fig. 2e). Whereas *Nanoarchaeum equitans*, one cultured representative in DPANN groups, integrates their cell membrane to its host^36^. This complex endomembrane structure may contribute to nutrition acquisition of *Nanoarchaeum equitans*^36^. However, this structure was not observed in TM7i; This indicates the difference between CPR and DPANN even though they adopt a similar lifestyle.

### “Luxury” type four pili gene is conserved in TM7’s extremely small genomes

We noticed that the highly streamlined genomes of TM7 contain highly conserved T4P gene clusters. As previously known, TM7, along with other CPR, generally has a small genome (less than 1 Mbp)^13^ and cell sizes. Therefore, the gene deletion of TM7 can be explained as an adaption to its symbiotic lifestyle: rely on its host for energy generation and bio-molecules synthesis. Based on whole-genome profiling of seven complete TM7 genomes including TM7i, we saw extremely reduced genomic repertoires with deficiencies in almost every metabolic pathway, including glycolysis, oxidative phosphorylation, and pentose phosphate pathways, which are seemed essential for bacterial physiology and lifestyle (Fig. 3a). Besides, the peptidoglycan biosynthesis pathway is complete in TM7’s genomes, indicating peptidoglycan synthesis is conducted in the cell cytoplasm. However, the T4P-assembly gene cluster, which is not vital to free-living bacteria, is highly conserved in these “small” genomes of parasitic TM7 (Fig. 3a). The T4P gene cluster contains pillin and pilC encode fimbrial proteins, pilB and pilT for pili elongation and retraction, and pilM. These proteins composed the molecular machine for a gram-positive bacterium to assemble or retract a type four pili^37, 38^. Multiple genomic alignment in all TM7 genomes also reveals T4P-related genes occurred in one of the major locally collinear blocks (LCB) (Fig. 3b). These results indicate T4P may play an indispensable role in TM7’s lifecycle.

**Figure 3.**
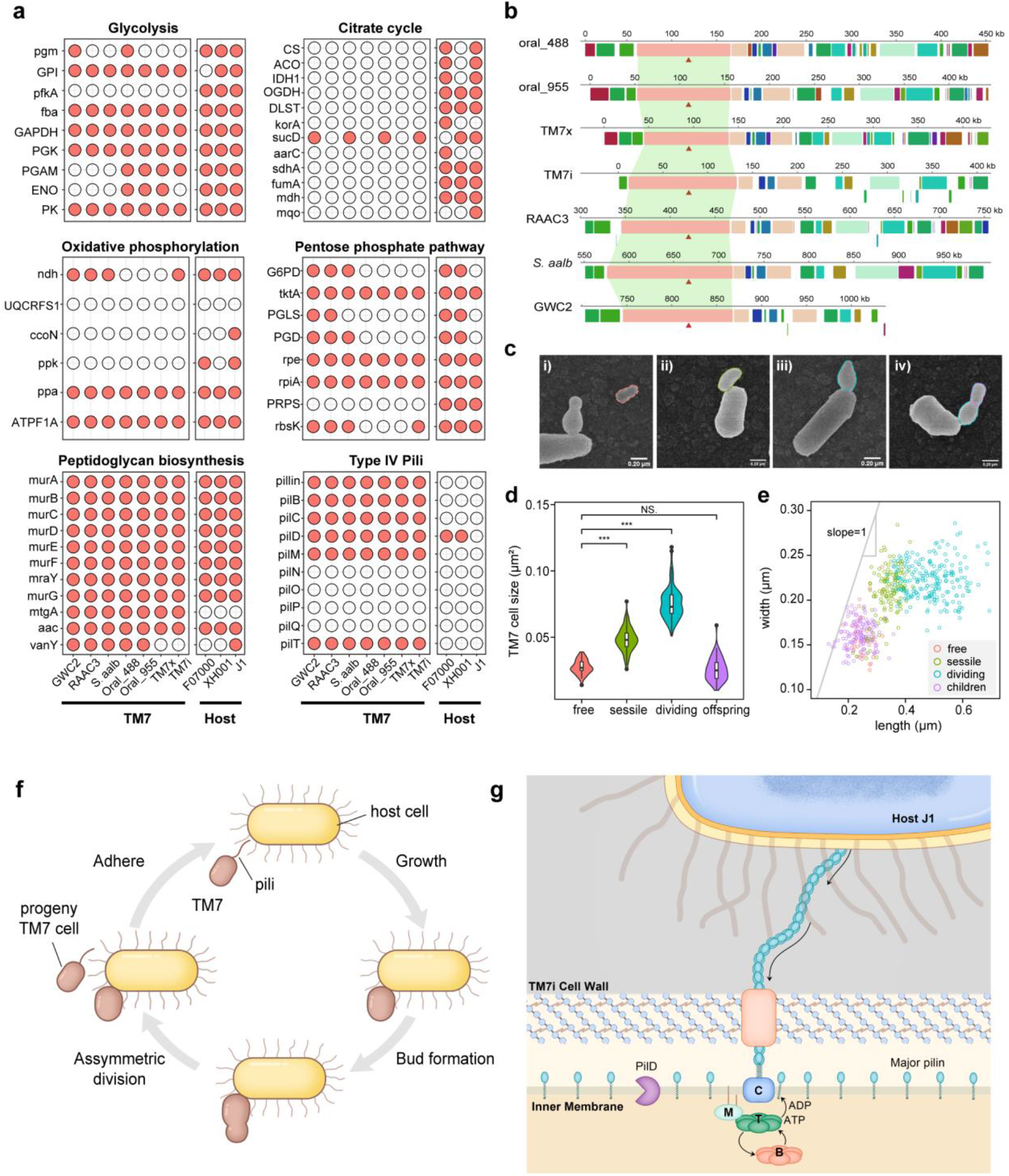
Predicted TM7i lifecycle based on comparative genomic analysis and microscopic observation. **a**, Metabolic potential of TM7 and their host indicates TM7 possess extremely stream-lined genomes and conserved T4P gene clusters. **b**, Locally Collinear Blocks matched in available TM7 genomes. Red triangle shows location of pilT, an ATPase for pili retraction. **c**, Differentiated TM7i cell volumes classified into four types: free living cells, newly adhered cells, dividing cells and offspring cells. scale bar=200nm. **d**, Violin plot of cell sizes quantitatively classifying the four cell types. **e**, Scatter plot of TM7i length and width classified in four cell types. **f**. Schematic illustration of the proposed lifecycle. **g**, Schematic diagram of predicted T4P assembly that initiate the cell-cell interaction between TM7i and its host J1 cell.

We envision that TM7 may rely on T4P to attach and drag forward itself to stick on their actinobacterial hosts tightly (Fig. 3f,g). First, as an epibiotic bacterium with serious metabolic deficiency, TM7 relies on close physical contact with its host for cell-cell interaction and metabolites exchange^14^. Secondly, T4P can functionally provide strong drag force^37^ and be employed by TM7 to make attachment to a host cell within reach, and be retracted to allow firm contact. Previous studies revealed that T4P is highly conserved in CPRs other than TM7^25^; this indicates CPRs may widely utilize the T4P to locate and attach to their hosts. Previous genomic-based study reveals Type Two Secretion System(T2SS), which includes a novel secreted array, is conserved in TM7 genomes^21^, which is also conserved in TM7i. As T2SS and T4P systems are evolutionarily related, we speculate that T4P and T2SS may play synergistic roles in the symbiotic interaction between TM7 and its host.

### Possible lifecycle of TM7

To investigate the enigmatic lifecycle of TM7, including its proliferation, nutrition acquisition, host selection, and parasitic mechanisms, we carried out microscopic observation and large-scale analysis of the TM7-J1 symbiont using SEM, TEM. More than 2000 TM7i cells were imaged using SEM, including free TM7i cells, sessile TM7i cells attached with J1 hosts, and budding TM7i cells with clear dividing sections (Fig. 3c-e). First, all microscopic images indicate that TM7i proliferates only after it is attached to the surface of a host J1 cell. Upon TM7i-J1 contacting interface, the tube-like structure can be seen in some SEM images (Supplementary Fig. 4a,b), which may help mass exchange between the host cell and the parasitic TM7 cell.

Based on SEM image analysis (Fig. 3d-e), significant morphological characteristics of TM7 cells were obtained, leading to our 4-stage lifecycle hypothesis: Stage (i): free TM7 cells have the smallest average width of 160 nm and length of 250 nm (n=41); Stage (ii): when a free TM7i cell attached on its host, it became a newly infecting cell, and its cell size gradually increases, with an increased average length of 337 nm, and cell area of 0.0487 μm^2^ (n=286); Stage (iii): budding stage: in this stage we found similar phenomenon as budding yeast, the attached TM7i cell generate a small daughter cell on its far end, and the daughter cells gradually expand its size. The lengths of a budding stage reached 693 nm; Stage (iv): binary fission stage: clear dividing section can be observed between the adhered cell and the daughter cell (Fig. 3c, Supplementary Fig. 4a). The daughter cells have a similar size as the free TM7i cell with average length of 238 nm (Fig. 3d,e). The SEM image analysis indicates that TM7i relies on its host for proliferation, and it has an uneven binary fission lifecycle with strict control of the cell size and shape (Fig. 3d-f). Interestingly, after being sessile upon their host, TM7i cell length and size began to grow synchronously with a slope close to 1; then, at the budding stage TM7i expands mainly its length (Fig. 3e). Combined with whole-genome profiling which suggests T4P-dependent interaction of TM7i with the host J1 cell, we propose the possible lifecycle of TM7i as shown in Fig. 3f: First, the free TM7i cell uses its type IV pili to attach to the host cell; second, the attached TM7i cell derive nutrition from the host cell for growth and proliferation, forming a smaller daughter TM7i cell on its far end; the matured daughter TM7i cell is released from the host-attached TM7i cell, starting its new lifecycle. TM7i cells of different stages can be distinguished based its size and morphology (Fig. 3c-f).

### The TM7i cell uses T4P for twitching motility

To confirm the existence and physiological function of T4P for free TM7i cells, we used Transmission electron microscopy (TEM) and structured illumination microscopy (SIM) to observe fresh separated TM7i cells from the filtrate of the TM7-J1 symbiont. The TEM images revealed that TM7i have typical type IV pili with a diameter of about 8 nm^37^ (Fig. 4a, Fig. 5c) and length of 0.2-2 μm, which is shorter than common gram-negative bacteria with a length of 2-5 μm^39, 40^. For live SIM imaging, TM7i cells are labeled with Alexa Fluor 555 probes, a non-specific fluorescence dye that labels all the protein upon and inside the cell. TM7i cells are constrained between a wetting layer between an agar block and a cover glass for continuous live SIM imaging at a frame rate of 10 seconds per frame. As shown in Fig. 4c, TM7i cells can dynamically extrude or retract their pili with time intervals usually less than 10s, which is typical for T4P^22^; this confirms again that the pilus filaments are T4P (Supplementary Video 3). Moreover, typical T4P-mediated twitching movements involving extension and retraction with attachment are visualized (Fig. 4c, Supplementary Video 1). The surface-attached pili first transitioned to a straight line, typically at a length of 0.361 μm, following by cell movements in the same direction of the straightened pili. The twitching movement direction is well aligned with the direction of the surface-attached pili. Among 73 twitching events measured, 91% of events have a shifted angle less than 5° (Fig. 4d). We observed the TM7i cells have twitching distances of around 0.335 μm per retraction (Fig. 4e). This is on average slightly shorter than the lengths of surface-attached pili measured for the same cells. And this type of twitching motility is in accordance with earlier study of T4P twitching study^41^.

**Figure 4.**
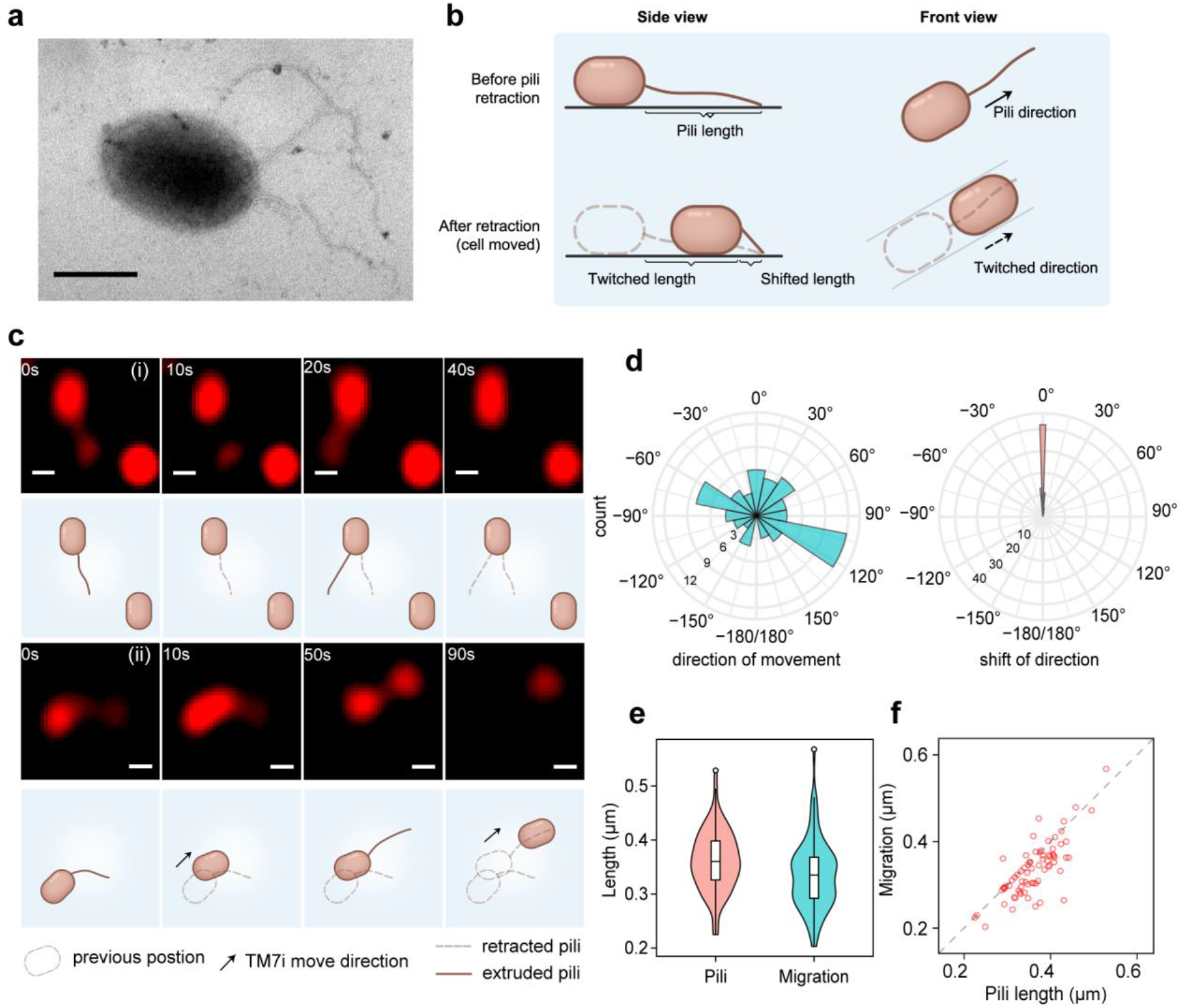
TM7i use T4P for typical twitching motility. **a**, A TM7 cell extruding T4P (scale bar=200 nm). **b**, Model exhibits TM7 T4P-mediated movement. **c**, Time-elapsed sequence shows TM7i T4P-mediated events: T4P extrusion and retraction (i, up shows the original fluorescence figures and bottom shows the illustration); continuous movement of TM7i (ii). (scale bar=200 nm) According to TM7i cell tracking (n=73), direction TM7i movements (**d**, left), the shift angle between T4P retraction and movement is shown at right (**d**, right). **e**, The length of T4P and TM7 cell movement. **f**, The relationship of the length of T4P-retraction and migration. The slope of the grey line is 1.

**Figure 5.**
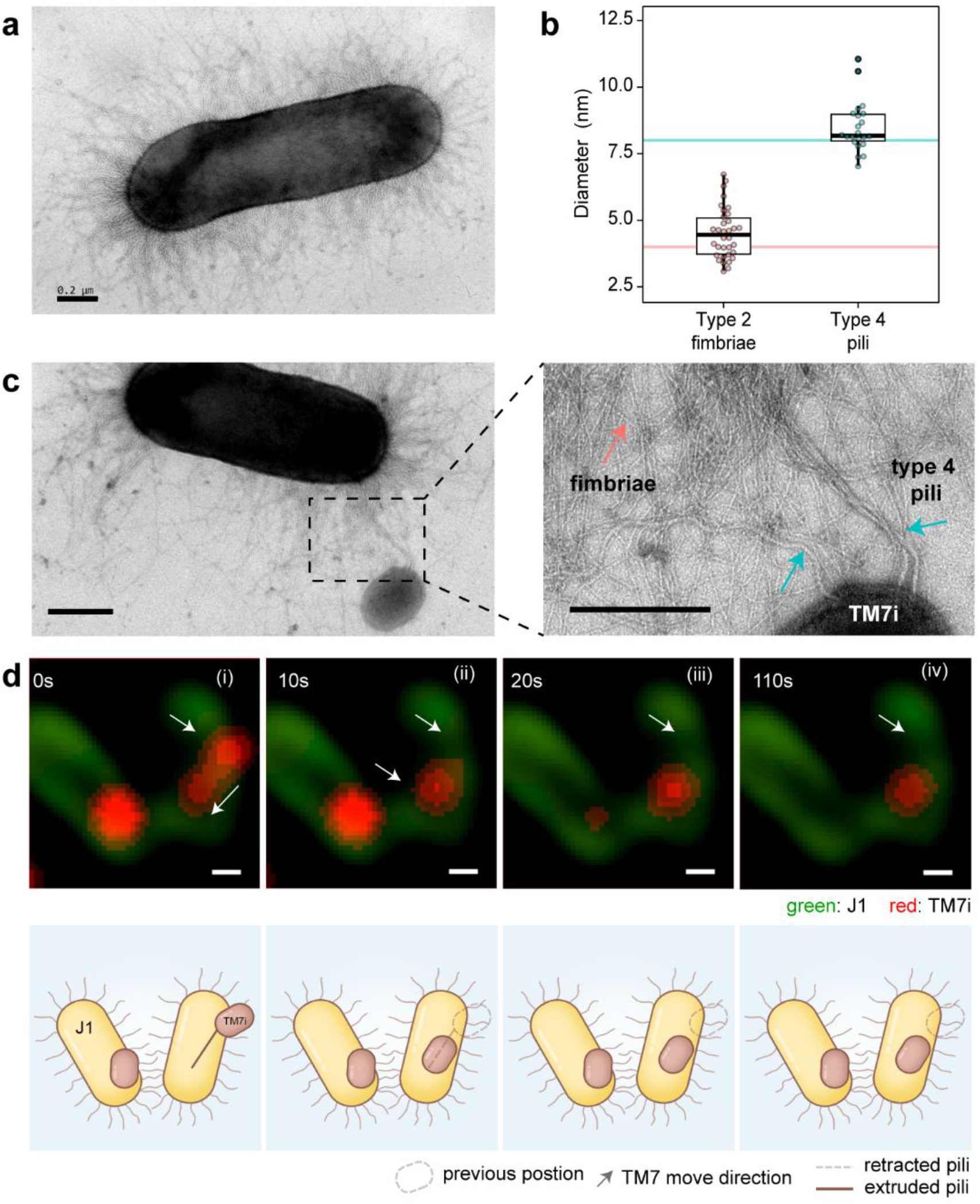
T4P helps TM7i attach to its host. **a**, A *Leucobacter aridocollis* J1 cell (the host) with hairy type 2 fimbriae. **b**, Comparison of T4P and fimbriae diameters of TM7i and J1, respectively. **c**, T4P of TM7i intertwined with fimbriae of the J1 cell. Red arrow indicates J1 type 2 fimbriae, blue indicates T4P of TM7i cell. **d**, Time series imaging reveals that TM7i uses T4P move towards J1. position of begin and end is marked by white arrows. The J1 and TM7i were labeled separately and their fresh mixture were imaged with 10s intervals. The bottom cartoon shows its illustration of the interaction process. (scale bar=200 nm)

### T4P possibly mediate TM7i-J1 interaction

To investigate the role of T4P in TM7i-J1 symbiotic interaction, we first used TEM to observe pure J1 cell as well as TM7i infected J1 cells. Its host, J1, has a type 2 fimbriae expression system according to its genome annotation. We preserved the type 2 fimbriae of the J1 cells with careful cell preparation. As shown in Fig. 5a, the J1 cell are covered with thin and hairy type 2 fimbriae with an average diameter of about 4 nm. The type 2 fimbriae of J1 can be distinguished from TM7i’s T4P according to its shorter lengths and thinner diameter (Fig. 4a, Fig. 5a,b). The TEM imaging captured the event of a free TM7i cell approaching a host J1 cell, with its T4P intertwined with type2 fimbriae of the J1 cell; this indicates that T4P might help the free TM7i cell to accomplish its parasitic lifecycle. As T4P enables twitching motility of free TM7i cells, it increases the opportunities of TM7i cells to approach and attach to its host cell and provides pulling force for dragging the TM7i cells toward the host to form an epibiotic symbiont. Moreover, the hairy type 2 fimbriae dramatically increase the surface area of the J1 cell, which makes locating of TM7i to J1 cell easier by increasing the probability of intertwinement between T4P and J1’s type 2 fimbriae.

To reveal the dynamic pili-related symbiotic interaction, we labeled TM7i and J1 separately with Alexa Fluor 488 probes and Alexa Fluor 555 probes, and mixed labeled TM7i and J1 cells to perform live-SIM imaging. The time-series SIM images revealed highly active motions of free TM7i cells (red) with many pili-dependent twitching events are captured, while J1 cells (green) remain motionless (Supplementary Video 1). In less than 3 minutes after mixing, a large portion of J1 cells are already infected by TM7i cells, which turned sessile as attached with its host. Interestingly, we observed some TM7i cells, moved toward uninfected J1 cells with T4P-mediated twitching motility, and became sessile when it was located on the J1 cell (Fig. 5, Supplementary Video 2). During the 110s time-lapsed imaging, the TM7i cell moved upon the J1 cell (0s-10s) and became immotile afterward (10s-110s), when the J1 cell remained still within the period. The results indicate that TM7i can actively locate and attach to its host with T4P-mediated motility.

## Discussion

Candidate phyla radiation (CPR) represents large evolutionary radiation whose members are largely uncultivated. TM7 is the first isolated obligate symbiotic CPR with representative small genomes and deficient biosynthetic pathways. The isolation of TM7x by streptomycin enrichment^14, 20^ shows its intriguing epibiotic lifestyle. Furthermore, new isolating methods such as targeted flow-cytometry^26^ significantly speed up the discovery of TM7 symbionts. However, knowledge about TM7 is inadequate to conclude its mysterious epibiotic lifestyle: mechanisms about its host recognition, symbiosis establishment, and proliferation are still unclear. Combined with the fact that Type IV pili are enriched in the genomes of CPR^25^ and its epibiotic lifestyle, it is easy to hypothesize that host-associated CPR microorganisms such as TM7 rely on T4P to accomplish their lifecycle. However, the lack of evidence remains to be an obstacle to a deeper understanding of CPR. Moreover, limited culture and limited knowledge about how CPR undergo their lifecycle hindered the advance of microbiology.

Our results first show that epicPCR can complement existing methods in CPR isolation by linking CPR to their hosts in the complex microbial community before cultivation efforts are exerted. As is firstly introduced to link the metabolic potential to phylogenetic markers in an environmental community^27^, epicPCR has also been used to reveal the virus-host interaction network in an estuarine environment^28^. In the present study, we use epicPCR to probe CPR sequences and fuse them with the 16S rRNA gene of its host co-encapsulated in picoliter emulsions, making it ideal for detecting the epibiotic landscope in a complex community. The epicPCR approach is more straightforward, efficient, and target-oriented than the existing methods. Meanwhile, by simply design and validate a set of concatenation PCR primers, epicPCR can efficiently screen a large number of samples and multiple symbiotic targets at the same time, revealing microbial epibiotic relationships at high throughput^28^.

We then developed a simple filtrate coculture workflow to isolate TM7 with host spectrum revealed by epicPCR and successfully obtained environmental-derived TM7i from a cicada-associated sample. The symbiosis of TM7i with *Leucobacter aridocollis* J1 shows an obligate epibiotic lifestyle similar to TM7x: an exclusive cell-cell attachment was observed in symbiosis. However, differences were also observed in the symbiosis: rather than being adverse to the growth of its host^12, 14, 42^, TM7i has a minor impact on J1’s growth status. Besides, little changes have been observed upon the shape and size of J1 cells during co-cultivation. This result can be explained by different characteristics of the habitat of symbionts and evolutionary biodiversity results in diversified adaption mechanisms. The newly isolated TM7i would give new insights into the mysterious TM7 phylum. Next, we profiled metabolic-related pathways and genes in both TM7 genomes and hosts’ genomes. We found that similar to recent opinions^25^, all representative TM7 have extremely streamlined genomes and lack major biosynthetic pathways. The small genome of TM7 is considered an adaptation of its epibiotic lifestyle, discarding genes essential to other free-living bacteria, such as key genes in the glycolysis pathway. However, a complete T4P expression cluster is found in all of the TM7 genomes. The comparison of conserved T4P and depleted “essential genes” indicates the importance of T4P for the parasitic lifestyle of TM7, which makes us wondered what role T4P might play in TM7i lifecycle, and how does TM7i accomplish its lifecycle.

Furthermore, we proposed the four-stage symbiotic lifecycle of TM7i with indications provided by quantitative analysis of electron microscopy and time-series SIM imaging. The lifecycle is possibly mediated by TM7i’s T4P, which is highly conserved in existing TM7 genomes, including our newly discovered TM7i. For the first time, we observed the dynamic motility toward the direction of T4P extrusion and retraction and the real-time union of TM7i-J1 symbiont, providing direct evidence for the critical roles of TM7i’s T4P in twitching motility as well as host recognition and infection. Meanwhile, we also observed the intertwinement of TM7i’s T4P with the hairy fimbriae of the J1 cell using thin-slice TEM. The type 2 fimbriae contribute to bacterial intrinsic adherence property^43^, But the function of the host’s type 2 fimbriae in TM7i-J1 symbiosis remains unknown. It will be worthwhile to investigate whether the length and density of fimbriae of the host contribute to the high host specificity of CPR groups^19^.

This study provides a first glimpse of the dynamic extrusion and retraction of T4P on the surface of nano-sized bacteria such as TM7. Using TM7i labeled with succinimidyl ester fluorescence probes, typical twitching motility of TM7i has been confirmed when nanometer cell displacement only happened in the direction of T4P reached out (Fig. 4d). The timescale of TM7i’s T4P mediated twitching movement is within tens of seconds. The length of pili and distance of cell displacement are highly interrelated (Fig. 4e). However, the SIM system we used is limited to a spatial resolution of 115 nm and a temporal resolution of 10s per frame, which is insufficient to visualize the 8-nm sized T4P clearly and 200-nm sized cells. The nonspecific succinimidyl ester fluorescence probes used in this study has been widely employed in labeling *Pseudomonas aeruginosa*^44^ and other bacteria^45^. Still, it provides limited fluorescence intensity with problematic photobleaching during long-term imaging. It’s still a challenge to selectively label T4P^39^ due to difficulties in genetically modifying TM7i. Other methods such as label-free interferometric scattering microscopy (isCAT) ^22^ are also limited due to TM7’s ultra-small cell size. Technical advances are required to improve the temporal and spatial resolution to characterize the TM7i-J1 interaction further.

### Conclusion

In summary, we report an epicPCR-directed approach for CPR isolation and successfully co-isolated TM7i with its host from a cicada-associated sample. The sequence information from epicPCR provides direct guidance to uncover the symbiotic network within the complex microbial community. Further, with supports from microscopic characterization and genomic analysis, our study shed light on the important role of T4P in TM7i’s lifecycle. For the first time, twitching motility and host attachment mediated by T4P was captured for a CPR nanobacteria with the help of super resolution imaging, showing how TM7i gets closer and attaches to its host cell. Our results also revealed the asymmetric budding division of TM7i cells. We believe that this epibiotic lifestyle is likely adapted widely by the CPR superphylum, such as uncultivated SR1, OP11, *etc*. Question arises, such as how TM7i transports nutrition and energy from its host, whether TM7 can be pure-cultured to be employed by other uses. These questions need to be further explored.

## Methods

### Sample collection and enrichment

Cicada exuviae were provided as traditional Chinese medicine from a pharmaceutical factory from Hangzhou, China. We weighed 0.5 g cicada exuviae and added it into 50 mL brain heart infusion (BHI) for a 2 days enrichment in a 28 °C incubation shaker with a speed of 180 rpm. PCR test with TM7-specific primers (F1-tm7-580F and 910R, Supplementary Table 1) was performed to determine if TM7 exists in enriched samples. PCR products were run on a 0.8% agarose gel. For samples with TM7-PCR signal continues to exist for 3 continuous passages, epicPCR (Emulsion, Paired Isolation and Concatenation PCR) and sequencing were performed as follows.

### Emulsion, Paired Isolation and Concatenation PCR

EpicPCR protocol was performed as previously described^27^ with some modifications listed below. The enriched samples were suspended in sterile water and vortexed. After SYTO9-labeling, the cell density is adjusted to 10∼20 million cells per 30 μL based on fluorescence microscopic cell counting and dilution. For fusion PCR, a primer set (Supplementary Table 1) was used to replicate a ∼300bp sequence from the TM7 genomes, with a universal primer end. This end then initialized the amplification of the host sequence, thus producing a ∼800 bp fragment. Amplified PCR product was then extracted using StarPrep DNA Fragment Purification Kit (Genestar, Beijing, China) according to manufacturer’s instructions. After extraction, nested PCR was performed using the fused DNA amplicons to produce a ∼500 bp internal sequence with adapters for Illumina MiSeq sequencing. Finally, purified amplicons were subjected to equimolar library pooling and paired-end sequencing on Illumina MiSeq PE300 (Illumina, San Diego, USA) according to the standard protocols (Majorbio, Shanghai, China).

For data analysis, The raw 16S rRNA gene sequencing reads were demultiplexed, quality-filtered by fastp (version 0.20.0)^46^ and merged by FLASH version 1.2.7)^47^. Samples were distinguished according to the barcode and primers; clean read was further processed using QIIME2^48^. A python script was introduced to separate amplicon sequences which have both the TM7-primer and universal-primer fragments. Furthermore, separated sequences were re-loaded to QIIME2 and to perform strain determination. ggplot2 and ggtree in R were used to visualize the abundance of fused sequences compared with the 16S-rRNA gene sequencing result^49, 50^.

### Bacterial isolation and growth

*Alcaligenes* sp. was isolated directly from cicada-enriched sample. *Leucobacter aridocollis* J1 was isolated in BHI agar added with streptomycin (100 μg/mL) and nalidixic acid (30 μg/mL) at a 28°C broth culture. *Leucobacter aridocollis* J1and *Alcaligenes sp*. were added with filtrate of enriched sample separately (500 μL filtrate in 50 mL culture broth). To identify if TM7 cells grew, PCR amplification for TM7 16S rRNA gene was performed with TM7-specific primer (Supplementary Table 2). Gel analysis and Sanger sequencing were performed to identify the target DNA product. For PCR-positive coculture (TM7-*Leucobacter* sp.), the same assay was performed for every 2d-passage to confirm a positive symbiosis.

Pure, un-infected culture of *Leucobacter aridocollis* J1 was used as the negative control; J1 broth culture was also added with culture filtrate, which contains TM7i cells, to test the impact of TM7i infection on J1 growth. For culture filtrate, a 2d TM7i-J1 culture was used to pass a 0.22 μm filter to enrich TM7i. The J1 broth culture in the BHI medium was diluted to 1/100 from a culture of OD600=0.4. Different volume of TM7i filtrate (0/1/10/100 μL) was then added into the J1 culture. And BHI medium was added to reach a volume of 300 μL. Each group has 3 biological and 3 technical replicates. The growth curve of TM7i and J1 was measured at 28 °C 200 rpm using Bioscreen (Oy Growth Curves Ab Ltd, Finland).

### Genome sequencing and assembly

The whole genome of TM7i and *Leucobacter aridocollis* J1 were sequenced using PacBio Sequel platform and Illumina NovaSeq PE150. For Library construction using PacBio Sequel platform, libraries for single-molecule real-time (SMRT) sequencing were constructed with an insert size of 10 kb using the SMRT bell TM Template kit (Pacific Biosciences, California, USA). For Library construction using the Illumina NovaSeq platform, A total amount of 1 μg DNA per sample was used as input material for the DNA sample preparations. Sequencing libraries were generated using NEBNext® Ultra(tm) DNA Library Prep Kit for Illumina (NEB, USA) following manufacturer’s recommendations and index codes were added to attribute sequences to each sample.

Preliminary assembly was performed with SMRT Link software (version v5.0.1). To ensure the accuracy of the subsequent analysis results, the low-quality reads were filtered (less than 500 bp) to obtain clean data; Using the automatic error correction function of SMRT portal, the long reads were selected (more than 6000bp) as the seed sequence, and the other shorter reads were aligned to the seed sequence by Blasr. The assembly was then corrected by the variant Caller module of the SMRT Link; the arrow algorithm was used to correct and count the variant sites in the preliminary assembly results. The assembly result was finally corrected with Illumina data. The corrected assembly result, which was used as the reference sequence, was blasted with Illumina data. Furthermore, the result was filtered with the base minimum mass value of 20, the minimum read depth of 4 and the maximum read depth of 1000. Cyclization of the assemblies were based on the overlap between the head and the tail.

### TM7 Pan-genomic analysis

For bacterial metabolic pathway analysis, all representative genomes were downloaded from NCBI, and annotated in KEGG by BlastKOALA^51^. Each K number annotated in genomes was classified into groups represented by EC number, which controls specific processes in a whole pathway. Multiple genome alignments were performed using progressive mauve with default parameters^52^.

### Electron microscopy

TM7i-J1 symbiont was passaged and cocultured for 24 hours and was prepared for electron microscopy procedures. Ultramicrotomy was employed in sectioning the symbiont cells for observation using TEM (Transmission electron microscope, JEM-1400, JEOL, Japan). Ultrathin sections of about 70 nm thickness were stained with 2% uranyl acetate and lead citrate. Negative stain was used to observe the intertwinement of pili-fimbriae of the symbiont under TEM. Scanning electron microscopy (SEM) was performed using HITACHI SU8018, before which Critical Point Drying (LEICA EM CPD300) and gold coating(E-1045) were employed. Images were captured at 5.0 kV.

### High-resolution fluorescence imaging of TM7i

Fluorescence probes (Macklin, Shanghai, China) of Succinimidyl esters linked with Alexa Fluor 555 or Alexa Fluor 488 were used to label surface proteins of TM7i and J1 cells separately as described^44, 45^. Free TM7 cells was isolated in the filtrate from a 2-day TM7i-J1 coculture by passing through a 0.45 μm filter (Jinteng, Tianjin, China). For both J1 and TM7i, 7000×g centrifugation for 10 min was used. The cell pellets were washed with 1 mL M9 medium for 3 times and resuspended in 0.5 mL M9 medium. We then added 40 μL of 1 mg/mL fluorescence probes and 35 μL of 1M sodium bicarbonate to raise the pH to 8.0∼8.5, and mildly shook the sample for 60 min at room temperature in the dark. After labeling, cells were centrifuged and washed twice with 1 mL M9 medium and resuspended in a certain volume of M9 medium (depends on cell number, typically 500 μL) to achieve an ideal concentration for imaging. For SIM imaging, 2 μL cell suspension of TM7i only or mixture of TM7i and J1 was added onto Nunc(tm) glass bottom dish (ThermoFisher, USA). Then, a 1% agarose pad of M9 medium was put onto the cell suspension. Fluorescence imaging was performed on the Super-Resolution Microscope System (N-SIMS, Nikon, Japan) using a 100× objective (Plan Fluo, NA 1.40, oil immersion) which can achieve a 115-nm spatial resolution. Figures were taken with an exposure time of 500 ms, interval of 10 s.

### Pili-related measurement

Measurement of pili was performed with ImageJ (NIH, USA). For pili length measurements in fluorescence studies, line length connecting center of TM7i cell and tip of pili was measured. For twitched length, line length between center of movement begins and end was measured. Shifted angle, which represents the difference of pili direction and twitched direction, was calculated. Diameters of T4P of TM7i and type 2 fimbriae of J1 were measured based on thickness of pili perpendicular to pili direction.

## Acknowledgment

We thank Professor Songnian Hu, Dr. Yingfeng Luo from the Institute of Microbiology Chinese Academy of Sciences for helpful discussion, and Dr. Chunli Li, Dr. Lei Su, and Miss Jingnan Liang for help in SEM, TEM and SIM imaging. This work was supported by National Natural Science Foundation of China (21822408, 91951103), National Key Research and Development Program of China (2018YFC0310703), International Ocean Resource Survey and Exploration (DY135-B-02), Center for Ocean Mega-Science, Chinese Academy of Sciences (KEXUE2019GZ05), and the Key Program of Frontier Sciences of the Chinese Academy of Sciences (QYZDB-SSW-SMC008).

## Data availability

The genomes of TM7i and *Leucobacter aridocollis* J1 have been deposited in NCBI GenBank under accession code CP075337 and CP075339, respectively. The raw reads of information generated from epicPCR were deposited into the NCBI Sequence Read Archive (SRA) database (BioSample Accession Number: SAMN19276902).

## Author contributions

W.D. and B.X. conceived and designed the project; B.X., J.W. and D.C. performed the research. B.X., W.D. and J.W. analyzed the data and wrote the draft; All authors contributed to writing of the paper.

## Additional information

**Supplementary information** is available for this paper.

## Competing interests

The authors declare no competing financial interests.

## Notes

### Competing Interest Statement

The authors have declared no competing interest.

