## Supplementary Information for "EpicPCR-Directed Cultivation of a *Candidatus* Saccharibacteria Symbiont Reveals a Type IV Pili-dependent Epibiotic Lifestyle"

**Supplementary Table 1.** primers used in TM7 detection and epicPCR.

| Primer | Sequence(5'to3') | Description |
|---|---|---|
| R1-F2-27F-910R | CTGAGCCAKGATCAAACTCTGTCCCCGTCAATTCCTTTATG | used in fusion PCR, epicPCR. |
| F1-tm7-580F | AYTGGGCGTAAAGAGTTGC | used in PCR to detect if TM7 exists and epicPCR |
| R2-519R | GWATTACCGCGGCKGCTG | used in fusion PCR, epicPCR |
| nest_806F | ATTAGAWACCCBNGTAGTCC | used in nested PCR, epicPCR |
| nest_341R | CTGSTGCVYCCCRTAGG | used in nested PCR, epicPCR |
| 910R | GTCCCCGTCAATTCCTTTATG | used in PCR to detect if TM7 exists. |

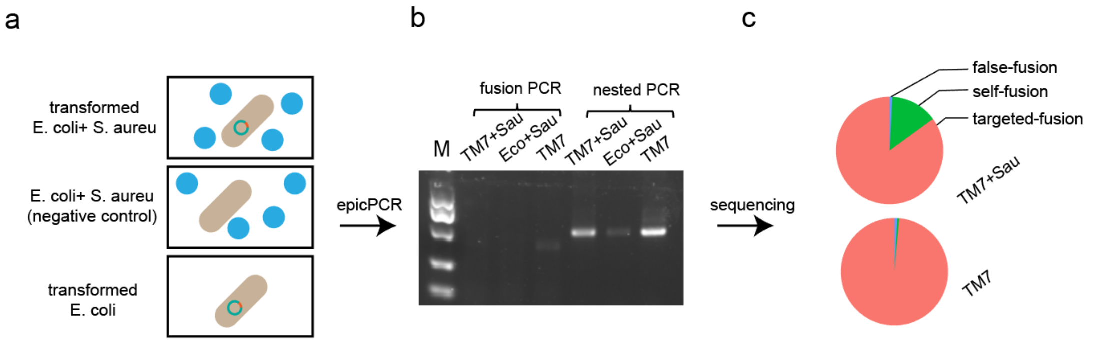


**Supplementary Figure 1.** Modified epicPCR system for symbiont detection is established and tested. (a) Sample used in epicPCR to test its capability for detecting microbial symbionts. TM7: Escherichia coli transomed with TM7 16s rRNA sequence; Eco: wild-type Escherichia coli; Sau: Staphylococcus aureus. When mixed, optical density ratio of 1:1 was referenced. (b) Gel analysis of epicPCR products. Because TM7-specific primer triggers the whole reaction, sample with no TM7 sequence had no amplification. (c) Amplicon sequencing result of TM7+Sau group and TM7 group. Since TM7 sequence was introduced into E. coli genome, targeted-fusion means transformed TM7 sequence fused with E. coli 16S rRNA gene; self-fusion indicates TM7 sequence fused with TM7 sequence; false-fusion indicates TM7 fused with Staphylococcus aureus sequence.

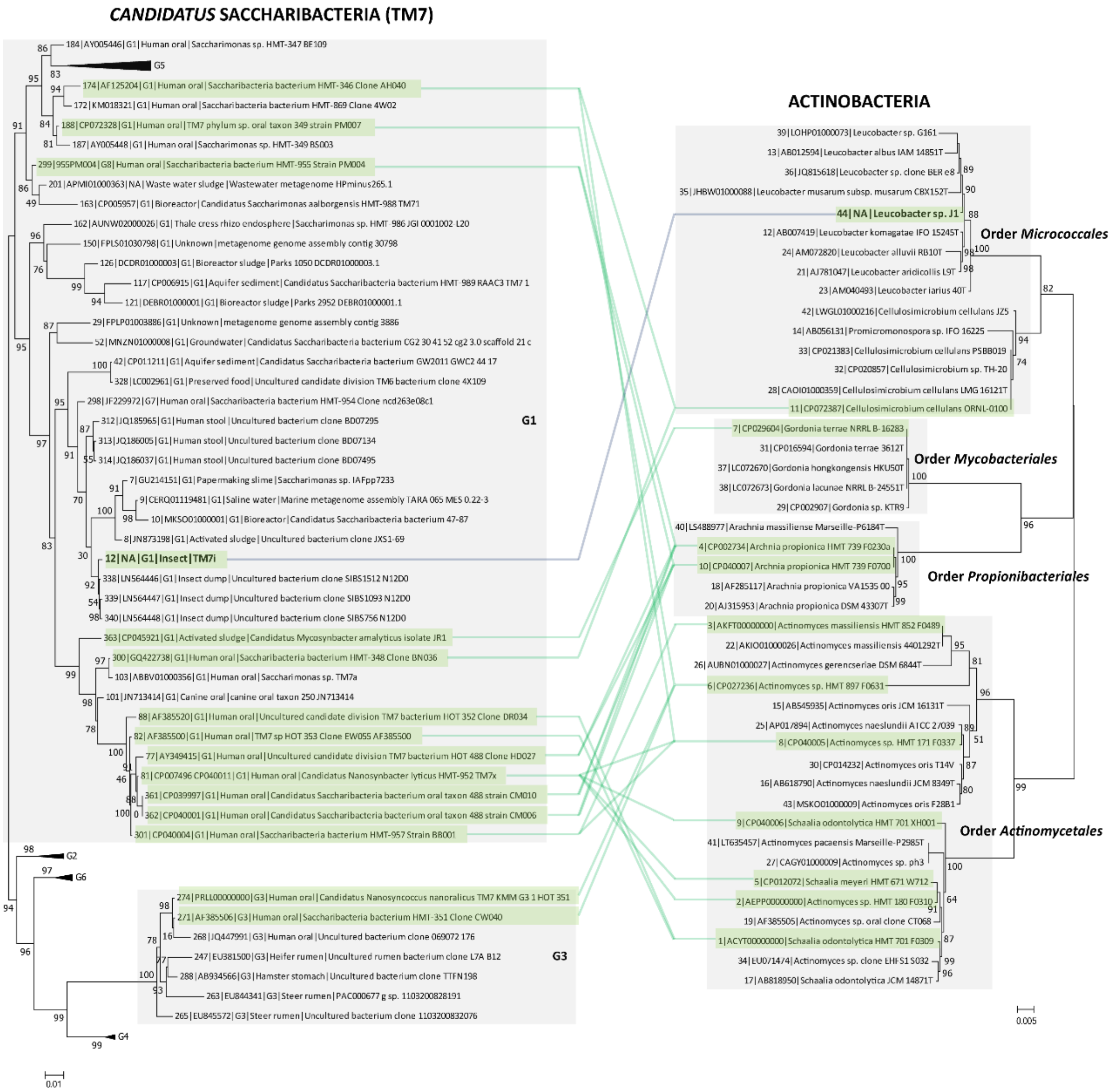


**Supplementary Figure 2.** Maximum-likelihood phylogenies with 16S rRNA gene sequences from the ARB/SILVA release 138.1 database showing the placement of strain TM7i within the G1 group of the phyla of *Candidatus* Saccharibacteria (TM7 group bacteria). The TM7 bacteria included in this tree were detected from human oral or open environments, identified by sequence accession numbers from GenBank/EMBL/DDBJ and associated with six operational groupings (G1-6). Lines connecting TM7 and Actinobacteria species indicate epibiont-host systems identified previously (green) and in this study (grey).

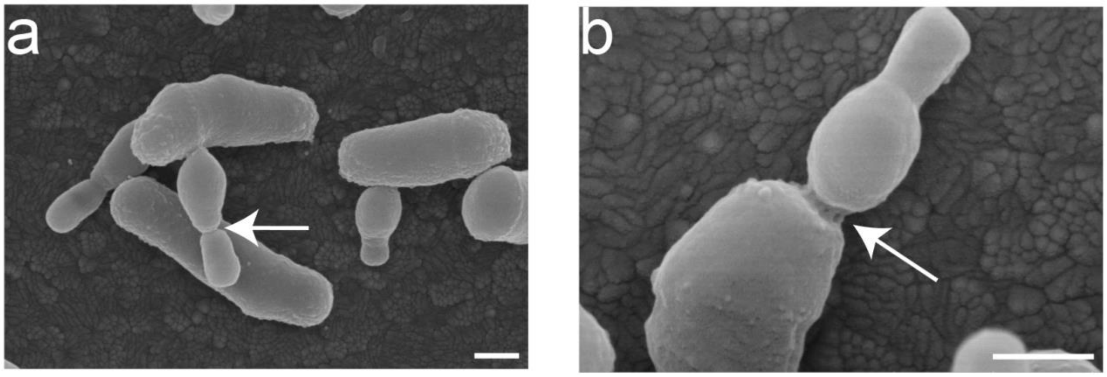


**Supplementary Figure 3.** uneven separation of TM7i and tube-like structure upon location. (a) SEM image showing TM7i proliferation. The offspring of TM7i is normally smaller than its parent. (b) SEM image showing tube-like structure between J1 and TM7i.
